# Galaxy-Kubernetes integration: scaling bioinformatics workflows in the cloud

**DOI:** 10.1101/488643

**Authors:** Pablo Moreno, Luca Pireddu, Pierrick Roger, Nuwan Goonasekera, Enis Afgan, Marius van den Beek, Sijin He, Anders Larsson, Daniel Schober, Christoph Ruttkies, David Johnson, Philippe Rocca-Serra, Ralf JM Weber, Björn Gruening, Reza M Salek, Namrata Kale, Yasset Perez-Riverol, Irene Papatheodorou, Ola Spjuth, Steffen Neumann

## Abstract

Making reproducible, auditable and scalable data-processing analysis workflows is an important challenge in the field of bioinformatics. Recently, software containers and cloud computing introduced a novel solution to address these challenges. They simplify software installation, management and reproducibility by packaging tools and their dependencies. In this work we implemented a cloud provider agnostic and scalable container orchestration setup for the popular Galaxy workflow environment. This solution enables Galaxy to run on and offload jobs to most cloud providers (e.g. Amazon Web Services, Google Cloud or OpenStack, among others) through the Kubernetes container orchestrator.

**Availability:** All code has been contributed to the Galaxy Project and is available (since Galaxy 17.05) at https://github.com/galaxyproject/ in the *galaxy* and *galaxy-kubernetes* repositories. https://public.phenomenal-h2020.eu/ is an example deployment.

**Suppl. Information:** Supplementary Files are available online.

**Contact:**, European Molecular Biology Laboratory, EMBL-EBI, Wellcome Trust Genome Campus, Hinxton, Cambridge, CB10 1SD, UK, Tel: +44-1223-494267, Fax: +44-1223-484696.

## 1 INTRODUCTION

The ever growing data volumes and data production rates in bioinformatics (Perez-Riverol et al. 2017) require the support of scalable and performant data processing architectures. Cloud computing has gained momentum, providing affordable, large-scale computing resources for high-throughput data processing. Bioinformatics has recently seen the adoption of this provisioning paradigm (Agapito *et al.*, 2017; Langmead and Nellore, 2018), through private clouds (e.g. local OpenStack (Sefraoui *et al.*, 2012)) and public cloud providers. However, most of these solutions are cloud specific, making difficult to reuse in more than one cloud.

Software containers have been recently adopted in bioinformatics– and also by the computing industry at large–as a way to distribute and manage software. Additionally, the development of container orchestrators enables the coordinated execution of a set of containers to host long-running services (e.g., database, web server and application server working together in separate containers) and batch jobs, acting as a clusters for containers. Kubernetes (kubernetes.io), the current *de-facto* container orchestrator solution, executes container-based work-loads in scalable infrastructure on many different cloud providers.

The Galaxy workflow environment (Afgan *et al.*, 2018) allows researchers to create, run and distribute bioinformatics workflows in a reproducible manner. Recent advancements in Galaxy simplified tool dependency resolutions through Bioconda (Grüning, Dale, *et al.*, 2018) and Biocontainers (da Veiga Leprevost *et al.*, 2017). These have paved the way for the use of container orchestration systems in the cloud with Galaxy.

Deploying software on cloud infrastructure normally requires specialised expertise that is foreign to most bioinformaticians. Moreover, each cloud provider implements different mechanisms, further complicating the matter. In the Galaxy context, CloudMan (Afgan *et al.*, 2012) was the first solution available to deploy Galaxy on Amazon Web Services (AWS), but support for other clouds requires extensive development efforts.

Here we introduce Kubernetes-based container orchestration and provisioning for Galaxy. Kubernetes acts as an abstraction layer between Galaxy and the different cloud providers, allowing Galaxy to run on every cloud provider that supports Kubernetes (>10 cloud providers currently). The solution described here combines ease of deployment, replicability of deployment, scalable computational performance, support for multiple cloud providers and use of containers to offload tools for distributed processing.

## 2 GALAXY-KUBERNETES ARCHITECHTURE

The Galaxy-Kubernetes setup is divided in three main concepts: (i) the Kubernetes Runner for Galaxy; (ii) the Galaxy container image with Kubernetes support; and (iii) packaging to automate the deployment of Galaxy in Kubernetes (Galaxy Helm chart). The Kubernetes Runner translates Galaxy jobs into Kubernetes jobs, specifying the container that needs to be executed for the Galaxy tools, and wrapping the commands that the job needs to execute for running them inside the container, once it is deployed to the Kubernetes cluster. The runner deals with the shared file system between Galaxy and the container running the tool (a requirement for this integration) (**Figure 1**). In addition, the runner handles user permissions, tool CPU/RAM requirements and the general management of the Kubernetes job lifecycle for Galaxy. The runner sends the job to Kubernetes and then enquires about the state of the job. This runner, after some minimal configuration of the appropriate plugin field and tools desired in the Galaxy job configuration file, is able to operate within any Kubernetes cluster that shares a filesystem with Galaxy.

**Figure 1:**
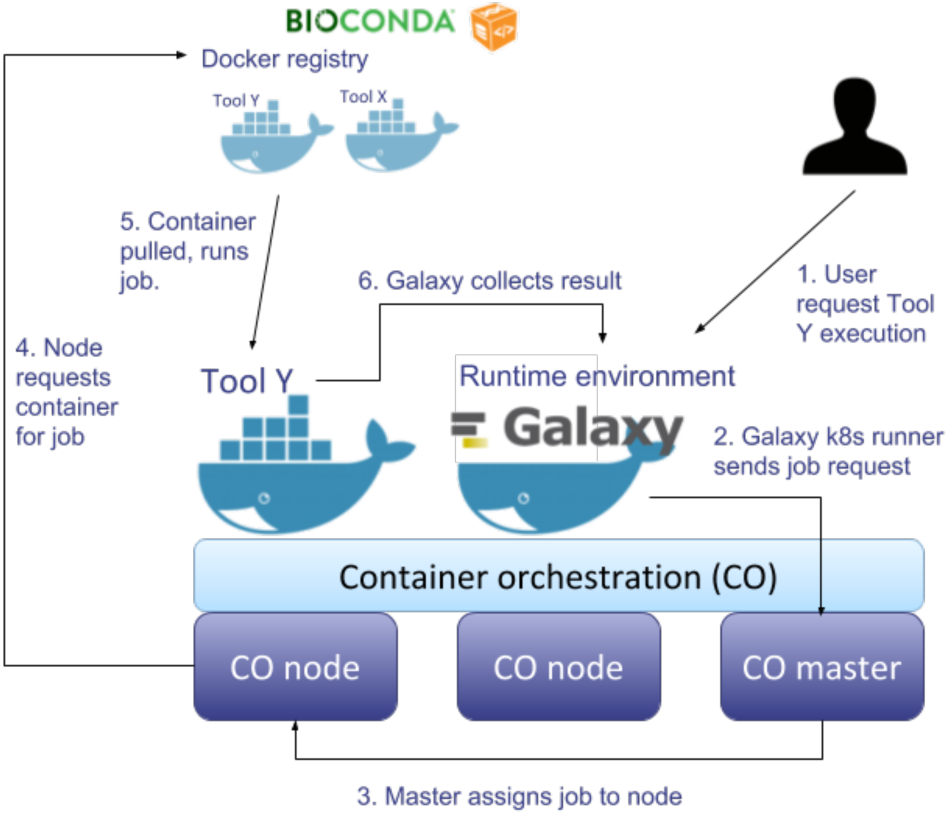
Galaxy running within Kubernetes as a container, and offloading jobs through containers to other nodes in the same Kubernetes cluster.

At this point, the user would still need to install Galaxy manually and plug it into a Kubernetes cluster. To make the integration process easier for developers and bioinformaticians, we improved the *docker-galaxy-stable* community Docker images (Grüning, Chilton, *et al.*, 2018), enabling the deployment of Galaxy through these composable containers on Kubernetes. This includes automated routines for runtime setup (using the Galaxy API (Sloggett *et al.*, 2013)) of users and workflows, installation of needed packages and additional testing of the container images, among others. Reusing the Galaxy community images reduces the burden of maintainability in the long run and improved the setup with many of the features that the community images have (e.g. web scalability, *sftp* access, etc). This standardisation permits the deployment of Galaxy instances that are complete replicates of each other on different cloud providers, maximizing interoperability and reproducibility.

Deployment of cloud-based solutions is a complex task that demands knowledge of both architecture (e.g. AWS, Google Cloud) and bioinformatics tools. We packaged the Galaxy setup for Kubernetes through Helm charts (helm.sh), which simplifies deployment on any Kubernetes cluster. Currently, the Galaxy Helm chart gives the administrator complete control of most of the configuration variables of Galaxy. This permits setting up different deployments (setups for different clouds, different Galaxy flavours/versions, testing environment, *etc*) as configuration files that can be version controlled (see examples, in the supplementary materials).

## 3 DISCUSSION AND FUTURE DIRECTIONS

We have implemented an approach to exploit encapsulation, packaging and container orchestration for Galaxy workflows, that simplifies cloud portability. This has been used extensively in the context of the PhenoMeNal H2020 Project (Peters *et al.*, 2018), in production at https://public.phenomenal-h2020.eu/ for the past three years (handling tens of thousands of jobs in the public instance). However, the set-up and methods are generic and can be easily applied to other omics, as done recently for instance on Single Cell RNA-Seq. Documentation linked on the supplementary materials shows how to use it.

## Supporting information

Frequently asked questions

Kubernetes runner details

## ACKNOWLEDGEMENTS

PM, LP, PRM, SH, AL, DS, CR, DJ, PRS, RW, RMS, NK, OS, and SN were supported by the European Commission’s Horizon 2020 program grant 65424 “PhenoMeNal”, and acknowledge the leadership of Christoph Steinbeck;

## CONTRIBUTIONS

Design and envisioned Galaxy-Kubernetes integration: PM; Code contributions Galaxy Kubernetes Runner: PM, LP, PR, MVB; Write Galaxy Helm Chart: PM, LP, NG, EA, SH, AL; Contribute to docker-galaxy stable: BG, PM, LP, SN; Manuscript writing: PM, LP, PRS; Galaxy Kubernetes testing: PM, LP, PR, CR, DS, DJ, PRS, RW, RM, YP, SN; Integration Kubenow: AL; Work package lead: PRS, OS, SN; Galaxy expertise guidance: BG; Project Manager Organize hackathons.: NK; Biocontainer interface: YP; Galaxy-Kubernetes Single Cell: PM, IP

