## Supplementary material for "Galaxy-Kubernetes integration: scaling bioinformatics workflows in the cloud": Frequently asked questions

### FAQ

#### Shared file system: does it impact minikube usage?

No, since Minikube runs as a single node Kubernetes cluster, the disk is by default available to both Galaxy and its jobs. So this setup can run on minikube without issues (besides the fact that you are by default limited in terms of resources).

#### Shared file system: Kubernetes on a cloud provider normally does not come with one, how do I get one?

There are many ways of doing this, and if you aim to run in production deciding how you will provision shared file system, and with how much resources, will be central to your ability to scale up Galaxy on Kubernetes. Some alternatives:

1. Pre-provisioned StorageClasses

Depending on your cloud provider and setup, it might be the case that you already have pre-set storage classes to some shared file systems (such as on demand GlusterFS, NFS or CephFS to name a few). If this the case you can directly make use of them by creating a persistent volume claim (PVC) that uses that particular storage class (note that it needs to be a ReadWriteMany compatible storage class), and then feeding that PVC to the Galaxy-Kubernetes setup, either through Helm or manually.

2. Provision shared filesystem on top of Kubernetes

Solutions like Heketi/Gluster-Kubernetes or Rook allow you to provision GlusterFS and CephFS on-demand respectively. This of course requires that you have sufficient privileges in your k8s cluster to be able to deploy these solutions on top of it.

3. Use a k8s provisioner that includes shared file system support

You can use solutions like Kubespray (GlusterFS supported only on OpenStack for the latter) or KubeNow (GlusterFS supported accross the board, but cluster has no high availability), which will provision a shared file system alongside Kubernetes. There are surely other provisioners that can produce something similar.

4. Manually provision or make shared file system available

You might have already and NFS server or alike setup within your institution. Provided that you have privileges to mount it and it is compatible with Kubernetes (there is a Persistent Volume that abstract it), then you should be able to use. There are also many Ansible/Puppet/Salt setups for deploying shared file systems out there.

#### Can I use cinder volumes in OpenStack as direct shared file system?

Unfortunately cinder volumes can be mounted to a single machine/VM, so they very much act like a USB disk that you plug to a single machine at a time. This means that you wouldn't be able to access them concurrently from different pods and/or kubernetes nodes.

#### How do I bake my own setup with my own tools?

When your setup doesn't require anything further than a release Galaxy version, then you need to:

- Create your own galaxy-init container with your tools. You can base your own galaxy-init container on the one from PhenoMeNal or this one from EBI Expression Atlas Group for Single Cell tertiary analysis. Basically, on your init container you will add:
  - Your tools, either as a set of tool ids (preferred) or directly adding the tool xml files.
  - Configuration files (see `config` dir in both example).
  - Ideally resource boundaries (RAM, CPU) per tool.
- Push the above container to a registry.
- Write your Helm config file so that your container, and any other injectable setting, is used to spin up your instance. Besides the example files on this repo, you can also check the EBI Expression Atlas Team example Helm config.
- Deploy on a Kubernetes cluster with Helm.

If however you require to use a development version of Galaxy, you might need to build first your own version of the docker-galaxy-stable container images using this script. Then your galaxy-init image needs to be based on the one created there.

#### Can I test deployments of Galaxy on Kubernetes within a CI like Travis?

Yes, we have an example at the EBI Expression Atlas Team Single Cell Galaxy setup.
