## Supplementary material for "Galaxy-Kubernetes integration: scaling bioinformatics workflows in the cloud": Kubernetes runner details

### Galaxy-Kubernetes Runner

If you are reading a PDF, the most up-to-date version of this document will be available here.

The Galaxy Kubernetes Runner allows Galaxy to offload containeraised workflows (in the form of docker containers) to a Kubernetes cluster that shares a file system with Galaxy. A shared filesystem is a necessary requirement for this runner to work. Galaxy can either be running outside of inside of the Kubernetes cluster to where jobs will be scheduled.

The Kubernetes Runner will translate a Galaxy Job into a Kubernetes (k8s) Job API object.

For detailed settings of the k8s Runner, please see that section in the `config/job_conf.xml.sample.advanced` file in your Galaxy installation. In this document we will explain the main features of the Kubernetes Runner.

#### Container resolution

Every job in k8s requires to be run inside the container. In Galaxy, a tool can be assigned a container via different methods:

1. By relying in the mulled automatic container resolution through bioconda packages. For this set variable `enable_beta_mulled_containers: true` in the `galaxy.yaml` configuration file (or through the corresponding environment variable. In this case you need to make sure that you are using only tools that have as dependency a bioconda package.
2. By specifying a container `destination` in the `config/job_conf.xml` file in Galaxy for the tool. This option can override option 1.
3. Using logic inside a dynamic destination (see for instance the example used in PhenoMeNal, particularly the dynamic destination code, the container assignment file and the job_conf.xml file.

The above resolution is done by code in Galaxy that is foreign to the k8s runner, the runner will simply assign the container resolved for the tool to the job being executed.

#### Resource settings

One of the key needs for running production workloads on top of a Galaxy-Kubernetes setup is that each tool has well defined memory and CPU requirements, specially for very heavy tools or tools that are normally massively distributed. When k8s executes a container with a workload, if no memory and CPU boundaries are set, the container orchestrator will simply allow any desired amount of resources to be consumed by the container. If the overall workload is close to the total capacity of the k8s cluster then this cluster will become unusuable and k8s might start to fail (as it control plane containers will struggle to access the needed resources).

Currently this can be handled either at the destination level in the job_conf.xml file, by setting the resources level allowed per destination, or through the use of dynamic destinations as done for example within PhenoMeNal (see dynamic destination code, the container assignment file and the job_conf.xml file

#### Shared file system mounting

As part of the configuration - either set by a helm deployment for Galaxy running inside k8s, or at `config/job_conf.xml - you will need to determine the name of the k8s Persistent Volume Claim (PVC). The runner will take care of mounting that same PVC inside the container that was resolved to run the job. In the automatic helm deployment everything is set so that all the files that Galaxy tools might need to execute a job are inside the PVC, however, if you are setting up Galaxy outside of k8s, care needs to be exercised here.

##### Accessing restricted privilege shared filesystem

On occassions you will want to use shared filesystems that are provisioned not only for the use of Galaxy and where you need to make use of certain user/group identifier. For those cases, the k8s Runner allows to set the k8s supplemental group id or the k8s fs group (through the pluign setup in the `config/job_conf.xml` file. These settings trickle down to every job sent. If you are using this outside of the Helm chart deployment make sure that the manually provided PVC is compatible with the group id used in either of this two settings.

#### Resilience

When designing cloud applications, there is always the need to consider what would happen if resources are not readily available when needed or if their access is interrupted (ie. a VM is restarted or lost). For this, the k8s Runner sends Galaxy Jobs to k8s Jobs setting the default pod retrial to 3 times (this is configurable in the pluging section). This means that if something foreign to the job goes wrong (ie. a disk is no longer available), then k8s Runner will wait until a 3 failure of the job to declare it failed. This has the downside that for very long jobs, it might take longer to detect that they failed (although this retrial number can be overriden at the tool level in Galaxy).

#### Pull policy for Jobs

In k8s, the pull policy dictaminates how agressive should the pull policy of container images be. A pull policy of `always` will make sure that containers manifests are always pulled on evey execution of a job, regardless of whether the k8s node executing that job has already the container image or not. `Always` might be useful for development environments. This setting is trickled down to each job executed. For more information on k8s pull policies, see this article.

#### k8s Namespaces

In principle, Galaxy can determine on which k8s namespace it will run, and this should trickle down adequatly to the k8s jobs executed. However, this hasn't been extensively tested. In principle, this would allow to easily run multiple Galaxy instances and their jobs concurrently on the same Kubernetes cluster.

#### Auditing jobs

Towards being able to audit what was run on the k8s cluster, the k8s Runner doesn't delete jobs once they have succesfully run, but simply re-scales them to zero. This is not currently configurable, but it would be a nice feauture to have in the future.

### Future plans

In the future, and funding allowing, we expect to add the following new features, among others to the Galaxy-Kubernetes runner:

- Support for Jobs using private containers.
- Support for Galaxy Interactive Environments.
- Ability to run without a shared file system.
- Improved instrumentation of jobs (for retrieval of detailed CPU and memory usage metrics).
- Support automated tool resources boundaries (CPU/RAM) through data available in the Galatic Radio Telescope
- Support for side car deployments to enable parallel functionality to tools.
- Integrations with Dask and Apache Spark deployments withing Kubernetes for tools that require it.
