## Supplementary File 1 for "Galaxy-Kubernetes integration: scaling bioinformatics workflows in the cloud"

### Galaxy Helm Charts

This repo contains Helm charts for easily deploying Galaxy on top of Kubernetes, in a number of scenarios, as described below.

If you are reading this from a PDF, the most up-to-date version of this document will be here.

#### Requirements

- Helm installed: Please follow official instructions from here.
- Access to a Kubernetes cluster (with shared file system accessible through a Persistent Volume or equivalent).
  - For development purposes or local tests, the local Minikube environment can be used. Install minikube following official instructions.
- kubectl cli: The command line argument for connection to a Kubernetes instance (remote cluster or local minikube). If not installed as part of Minikube steps, follow ONLY the installation steps (not the configuration ones) from here.

#### Minikube

If using minikube, you need to make sure that it is running. If you just installed it, you need to execute `minikube start`. In general you can check the status of minikube through `minikube status`.

#### First time installation

If using helm for the first time, you will need to initialize the helm on the cluster and add the helm repo to the local helm directories:

```
$ helm init
$ helm list # call a few times until no error is shown, this is to wait for the tiller pod from helm to be running on the cluster.
$ helm repo add galaxy-helm-repo https://pcm32.github.io/galaxy-helm-charts
```

if you have done this once in the past, you might need, from time to time, to update the local repo, by doing:

```
$ helm repo update
```

#### `galaxy-stable` chart documentation

The main Galaxy chart is `galaxy-stable`, which is designed to run Galaxy using the docker-galaxy-stable compose container images. This setup follows the Galaxy recomendations for production setups. It will spin up a Galaxy container, and sftp server (ProFTPD, for data uploads) container and a PostgreSQL relational database container (used by Galaxy). The folder example-configs has helm configurations that can be used to spin different Galaxy setups.

##### Deployment example 1: PhenoMeNal - Galaxy 18.01 - Minikube

For instance, to spin up the PhenoMeNal Galaxy setup with metabolomics tools, you can execute (assuming the minikube case for testing and that you have checked out this repo or retrieved the example_configs directory):

```
helm install -f example_configs/galaxy-stable-phenomenal-18.01-minikube.yaml galaxy-helm-repo/galaxy-stable
```

After a few minutes, invoking `kubectl get pods` should show something like:

```
NAME                                                READY     STATUS    RESTARTS   AGE
impressive-kudu-galaxy-stable-74fd5cc5-qg5z4            1/1       Running   0          4m
impressive-kudu-galaxy-stable-proftpd-df446777b-wpdz9   1/1       Running   0          4m
impressive-kudu-postgresql-556744cfbd-5hjph             1/1       Running   0          4m
```

then you can check the ip of your minikube machine through:

```
$ minikube ip
192.168.64.4
```

in the case of this example then, Galaxy is be available at http://192.168.64.4:30700 on your local machine. To stop the instance, one would use the name of the helm deployment (can be obtained with `helm list`) to stop it. In the case of this example, this would be done with:

```
helm delete impressive-kudu
```

Minikube on its default setting might not expose sufficient resources to actually run tools (as the tools might be requesting more CPU/RAM than that allocated to minikube). To allocate more resources to minikube, provided your machine has them, you can start it with:

```
minikube start --cpus <number-of-cpus> --memory <memory-in-megabytes>
```

#### Available Chart variables

The following table includes all the variables that can be set up using the galaxy-stable helm chart to configure a deployment.

| Parameter | Description | Default |
|---|---|---|
| export_dir | Export directory for Galaxy compose | /export |
| init.image.repository | Repository for the docker image: <server>/<owner>/<image-name> for Galaxy init. | pcm32/galaxy-stable-k8s-init |
| init.image.tag | Image tag for Galaxy init image. | pcm32/galaxy-stable-k8s |
| init.image.pullPolicy | Pull policy for the Galaxy init image | IfNotPresent |
| init.resources | k8s resources map for the init process | |
| image.repository | Repository for the docker image: <server>/<owner>/<image-name> for Galaxy main process. | pcm32/galaxy-stable-k8s |
| image.tag | Image tag for Galaxy image. | latest |
| image.pullPolicy | Pull policy for the Galaxy image. | IfNotPresent |
| tools.destination | Directory where tools are stored, possibly not needed and should be removed. | /export/tools |
| k8s.supp_groups | Kubernetes supplemental group (this is probably a list), used for writing with adequate privileges to certain shared file systems | |
| k8s.fs_group | Kubernetes file system group (this is probably a list), used for writing with adequate privileges to certain shared file systems | |
| admin.email | Admin email to setup Galaxy with. | |
| admin.password | Admin password to setup Galaxy with. | |
| admin.api_key | Admin api_key to setup Galaxy with. | |
| admin.username | Admin username to setup Galaxy with. | |
| admin.allow_user_creation | Configures allow_user_creation Galaxy config environment variable. | "True" |
| galaxy_conf.* | Replace * for any variable name available in the galaxy.ini or galaxy.yaml main configuration file. Some examples below. | |
| galaxy_conf.brand | Branding text displayed on Galaxy | k8s |
| galaxy_conf.smtp_server | SMTP server for Galaxy password reset functionality | |
| galaxy_conf.smtp_username | SMTP username for Galaxy password reset functionality | |
| galaxy_conf.smtp_password | SMTP password for Galaxy password reset functionality | |
| galaxy_conf.email_from | SMTP email_from for Galaxy password reset functionality | |
| galaxy_conf.smtp_ssl | SMTP ssl for Galaxy password reset functionality | |
| galaxy_conf.url | Incoming URL label for Galaxy password reset functionality, shown on reset email to identify instance. | |
| galaxy_conf.allow_user_deletion | Allows the admin to delete users | |
| galaxy_conf.allow_user_creation | Allows the admin to delete users | |

| | | |
|---|---|---|
| galaxy_conf.containers_resolvers_config_file | Config file path for resolving containers | "/export/config/container_resolvers_conf.xml" |
| galaxy_conf.ftp_upload_site | Incoming URL for sftp uploads. | |
| service.port | Internal port where the Galaxy container serves content. | 80 |
| service.nodePortExposed | Internal port where the Galaxy container serves content. | 30700 |
| service.name | Name to use for the k8s service exposing Galaxy | galaxy-svc |
| service.type | Type of k8s service for Galaxy | ClusterIP |
| persistence.enabled | Whether to create or not a PVC for Galaxy, defaults to true. | true |
| persistence.existingClaim | Name of an existing read-write-many PVC to use. If it is given, no PVC is created. | |
| persistence.name | Name for the PVC that Galaxy and scheduled jobs will use. | galaxy-pvc |
| persistence.size | Amount of this that the PVC requests, such as "15Gi" | 15Gi |
| persistence.subPath | Subpath within the PV where the PVC should reside. | |
| persistence.minikube.enabled | Whether to create a Persistent Volume in minikube or not. | true |
| persistence.minikube.hostPath | Path in the minikube VM for galaxy data directory (where PV gets created). | |
| ingress.self_managed | Whether to use an external ingress controller or the chart's provided one, when using ingresses. | false |
| ingress.enabled | Whether to enable ingress or not... seems redundant, should be fixed. | false |
| ingress.hosts | Hostname array to construct ingress URLs to respond to (hostname.domain). | galaxy |
| ingress.domain | Domain to construct ingress URL to respond to (hostname.domain). | local |
| ingress.path | URL path for Galaxy | |
| ingress.annotations | | |
| ingress.tls | | |
| resources | Resources requests and limits (k8s) for the Galaxy container when running. | |
| postgresql.enabled | Whether Galaxy should use postgres or not. | true |
| postgresql.postgresPassword | Password to use for postgres setup | change_me |
| postgresql.postgresUser | User for postgresql | |
| postgresql.postgresDatabase | Name for the galaxy database | |
| postgresql.persistence.existingClaim | Name of the Persistent Volume Claim to use with postgres, by default the same as Galaxy | galaxy-pvc |
| postgresql.persistence.subPath | A subpath in the PV where the postgres mount will be done | |
| postgresql.fullname | ?? | |
| legacy.pre_k8s_16 | Whether we are running on a Kubernetes setup below 1.6 | false |
| rbac.enabled | Whether RBAC setups for the chart should be activated. | false |
| proftpd.enabled | Use proftpd or not | true |
| proftpd.replicaCount | Number of instances for proftpd. It is not clear whether >1 would work due to keys generation. | 1 |
| proftpd.image.repository | docker image for proftpd | |
| proftpd.image.tag | tag for the proftpd image set above | |
| proftpd.passive_port.low | Passive port minimum for proftpd | |
| proftpd.passive_port.high | Passive port maximum for proftpd | |
| proftpd.use_sftp | If set to true, use sftp instead of ftp | "true" |
| proftpd.service.name | Name to be given for the proftpd k8s service | |
| proftpd.service.type | Type of service for proftpd | ClusterIP |

| | | |
|---|---|---|
| proftpd.service.nodePortExposed | Port opened on k8s nodes for exposing proftpd, when using service type NodePort | 30722 |
| proftpd.generate_ssh_key | Whether to generate the ssh key for sftp access | "false" |

### Funding

Most of this work, including the Galaxy-Kubernetes integration, has been contributed to the community thanks to the funding of the European Comission (EC), through the PhenoMeNal H2020 Project, grant agreement number 654241.
